# The devil in citizen science data: observation processes invalidate the causal inference that photovoltaic policy reduces bird diversity

**DOI:** 10.64898/2026.08.25.746390

**Authors:** Yangkang Chen, Wenyuan Zhang, Heng-Xing Zou, Xu Shi, Yang Liu

**Author notes:** Corresponding author (Y.C.). Code and data are available at https://github.com/chenyangkang/reanalysis-Science-PSI-ShannonBD.

## Abstract

Citizen science data are increasingly used to infer biodiversity change, but causal claims based on such data are credible only if sampling effort and its temporal shifts are explicitly modeled. Zhang et al. (*1*) used citizen science data to conclude that greater photovoltaic policy stringency, measured using the photovoltaic policy stringency index (PSI), reduced county-level bird diversity in China. We reproduced their fixed effects and instrumental variable estimates. However, the observed Shannon diversity derived from pooled citizen science records reflects both bird communities and sampling effort, which the authors’ controls do not adequately capture. Accounting for observer count changed the reported statistically significant 2.10% decline in Shannon index to a nonsignificant 0.58% increase (*P* = 0.288) per one-standard-deviation increase in PSI, and rendered the instrumental variable estimate statistically indistinguishable from zero (*P* = 0.912). Yet observer count is only one of many sources of sampling bias. PSI was also associated with multiple dimensions of sampling effort, consistent with sampling effort acting as a potential mediator in the PSI–diversity chain. The sampling domain also shifted markedly from 2014 to 2023: recorded county-months increased almost 24- fold, median observer count rose from one to three, and zero-duration records declined from 57.2% to 0.17%. Without adequate adjustment, these shifts confound estimates of temporal change in observed bird diversity. Beyond its inadequate treatment of sampling effort, the original study also misinterpreted its statistical results. Although the reported *R^2^* values are high, they are dominated by county and year-month fixed effects, with PSI contributing a partial *R^2^* of only 0.048% on observed Shannon index. The PSI–photovoltaic-area correlation is also weak (*r* = 0.0414) and vanishes after accounting for fixed effects (*P* = 0.977). Furthermore, the released bird observation data contain many erroneous outliers, raising significant concerns about insufficiently rigorous data preprocessing and quality control. These results show that the released data cannot properly distinguish ecological change from sampling effort change. Robust inference from citizen science data requires checklist-level effort metadata, explicit correction for spatiotemporal sampling shifts, and close collaboration among researchers with complementary methodological and ecological expertise.

## Background

Assessing whether policies that accelerate solar deployment have unintended consequences for biodiversity is an important and timely research question. Zhang et al. (*1*) sought to address this question at a national scale by assembling an unbalanced county-month bird observation summary comprising 46,528 observations from 2,344 Chinese counties between 2014 and 2023. They calculated the Shannon diversity index of bird communities (ShannonBD) from species and abundance records obtained from the China Birdwatching Record Center (China Bird Report; https://www.birdreport.cn), a citizen science (participatory science) platform, and constructed a photovoltaic policy stringency index (PSI) by scoring official policy documents according to their administrative force and weighting policies issued at the provincial, municipal, and county levels. They estimated the association between PSI and bird diversity using county and year-month fixed effects regressions with environmental, socioeconomic, land-cover, and birdwatching duration controls. Their fully controlled model indicated that a one standard deviation (1 SD) increase in PSI was associated with a 2.10% reduction in Shannon diversity (*P* < 0.01). They further used an instrumental variable (IV) design based on historical sunshine and city-level climate-policy uncertainty, together with propensity score matching and other robustness analyses, to support a causal interpretation. Heterogeneity and mechanism analyses were then used to argue that stronger photovoltaic policy disproportionately reduced diversity in wealthier and non-desert counties and among certain species groups, primarily through land conversion, vegetation homogenization, and what the authors termed “inferior greening.”

Here, we test whether the reported PSI-diversity relationship can be separated from the observation process. We reproduce the original estimates, examine whether PSI predicts observation effort, and re-estimate the principal models with additional sampling controls. Citizen science records are valuable for large-scale ecological research, but unlike protocol-based surveys, their coverage and detection processes are determined by voluntary participation (*2–4*). Observer number and expertise, checklist completeness, survey duration and distance, sampling protocol, habitat accessibility, hotspot selection, and reporting practices can all affect recorded species richness, abundance, and Shannon diversity without any underlying ecological change (*4*). Moreover, uneven changes in platform adoption, observer composition, observer behavior, and reporting quality within counties can produce temporal sampling domain shifts that common year-month fixed effects do not remove (*5–7*). We reproduced the study’s fixed effect models and IV estimates, examined its statistical interpretations and the PSI–photovoltaic-area (PV area) relationship, evaluated the internal consistency of the released bird and sampling effort variables, tested whether PSI predicts sampling effort, and re-estimated the principal models with additional sampling effort controls.

## Results

Zhang et al. (*1*) reported that stronger photovoltaic policy, quantified by PSI, reduces county-level bird diversity in China. We reproduce their fixed effects and instrumental variable estimates but find that the causal interpretation is not supported by the released data.

### Sampling effort was not adequately controlled for

The observed Shannon diversity index is influenced by the observation process, which the authors may not have adequately accounted for (see the next section for explanation). Consistent with this concern, we found that **PSI is associated with the sampling process itself**: it predicts whether a county-month appears in the dataset, the number of observers, and total observation time. Although observer counts increased nationally over the study period, after accounting for common year-month trends, a one-standard-deviation within-county increase in PSI was associated with a 3.24% lower probability of sample inclusion, 12.5% fewer observers, and 20.7% less total observation time on the log scale. One plausible sampling-mediated mechanism is that photovoltaic development or other local changes associated with PSI made certain sites less accessible or attractive to birdwatchers. Reduced participation and coverage would lower species detections, observed richness, and observed Shannon diversity even without a decline in the underlying bird community. Consistent with this mechanism, observer count was strongly correlated with richness (*r* = 0.801, *P* < 0.001, Fig. 2) and Shannon diversity (*r* = 0.490, *P* < 0.001, Fig. 2), and all sampling effort variables increased throughout the years. Alternatively, the estimated PSI values might be inaccurate, since it is an index scored by the authors using the photovoltaic-policy documents from government websites. We cannot validate the PSI score since those documents are not available and the validity of the method being unverified.

The link between PSI and sampling effort matters substantively to the main results. **Adding observer count changes the coefficient of PSI on ShannonBD from negative and significant (-2.1%, *β*= -0.0125, *P* = 0.0006) to small, positive, and nonsignificant (+0.58%, *β*= 0.0035, *P* = 0.288, Fig. 1, Fig. 3)**. Similarly, the released IV model estimates a significant negative effect of PSI on Shannon diversity (*β*= -0.126, *P* < 0.001), but the estimate becomes essentially zero after controlling for observer count (*β*= 0.003, *P* = 0.912). The IV does not remove this sensitivity. Adding observer count changes the estimate from *β*= -0.12637, *P* < 0.01 to *β*= +0.00299, *P* = 0.912. Interestingly, the instrument itself also predicts observer count (-0.00656, *P* < 0.001), demonstrating that the identifying variation is associated with observation intensity. Because observer count strongly influences observed Shannon diversity, an instrument that predicts observer count is unlikely to satisfy the exclusion restriction. The IV estimate therefore cannot distinguish policy-related changes in sampling from changes in the underlying bird community unless observer-mediated sampling pathways are ruled out. The IV analysis therefore does not rescue the claimed ecological causal interpretation. These results reveal the observation process pathway, that the study failed to account for, can generate the reported PSI–diversity relationship.

**Figure 1.**
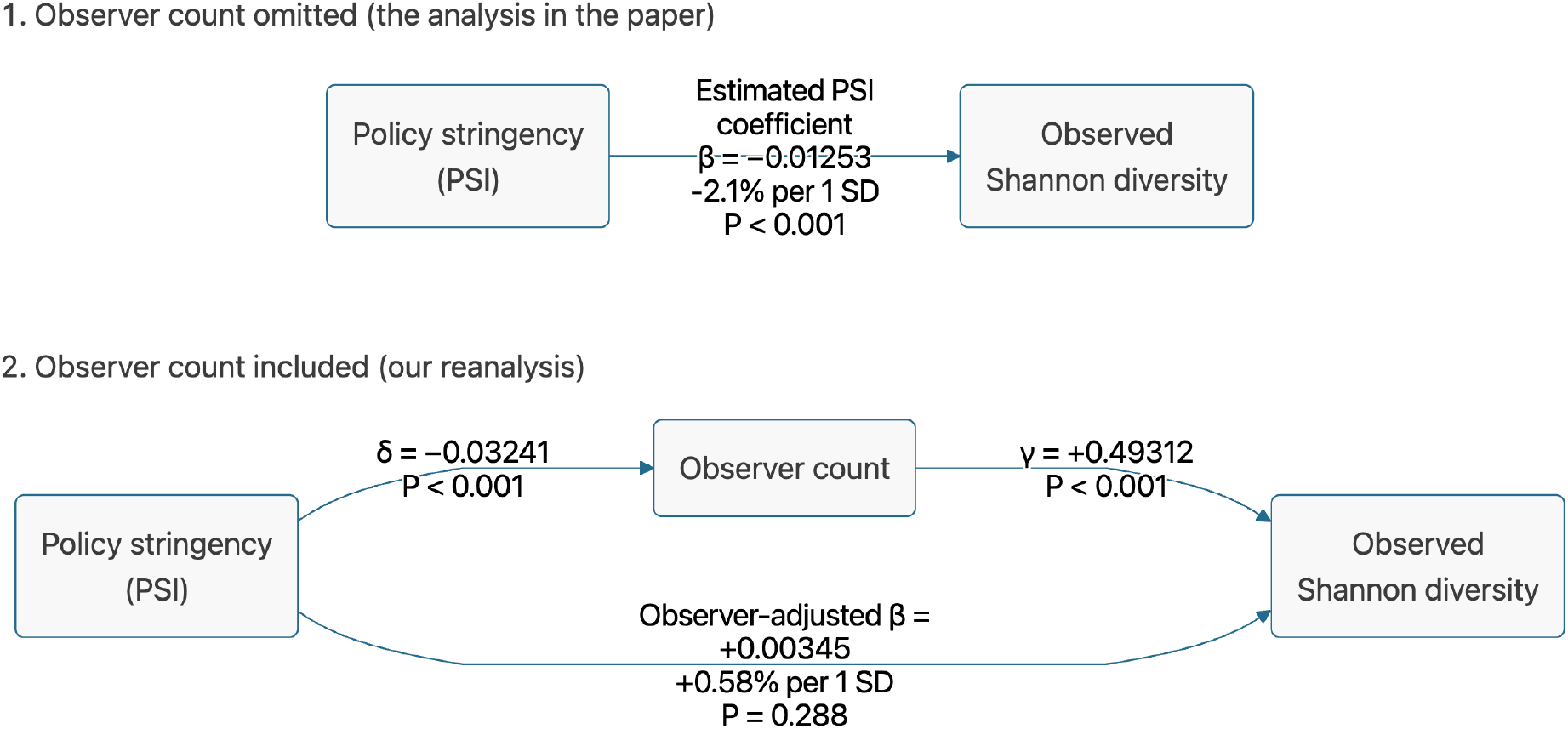
An alternative causal diagram of PSI and ShannonBD. **(upper panel)** Reproducing the study’s specification without observer count yields a negative PSI coefficient (*β*= -0.01253, *P* = 0.0006), corresponding to a 2.10% decrease in observed Shannon diversity per one-standard-deviation increase in PSI. **(lower panel)** After including log-transformed observer count, PSI is associated with fewer observers (*δ*= -0.03241), and observer count is positively associated with observed Shannon diversity (*γ*= 0.49312); the PSI coefficient becomes small, positive, and nonsignificant (*β*= 0.00345, *P* = 0.288), corresponding to a 0.58% increase per standard deviation. All models include the study’s original controls, county and year-month fixed effects, and county-clustered standard errors (*N* = 46,371). Arrows represent conditional regression associations, not established causal effects.

**Figure 2.**
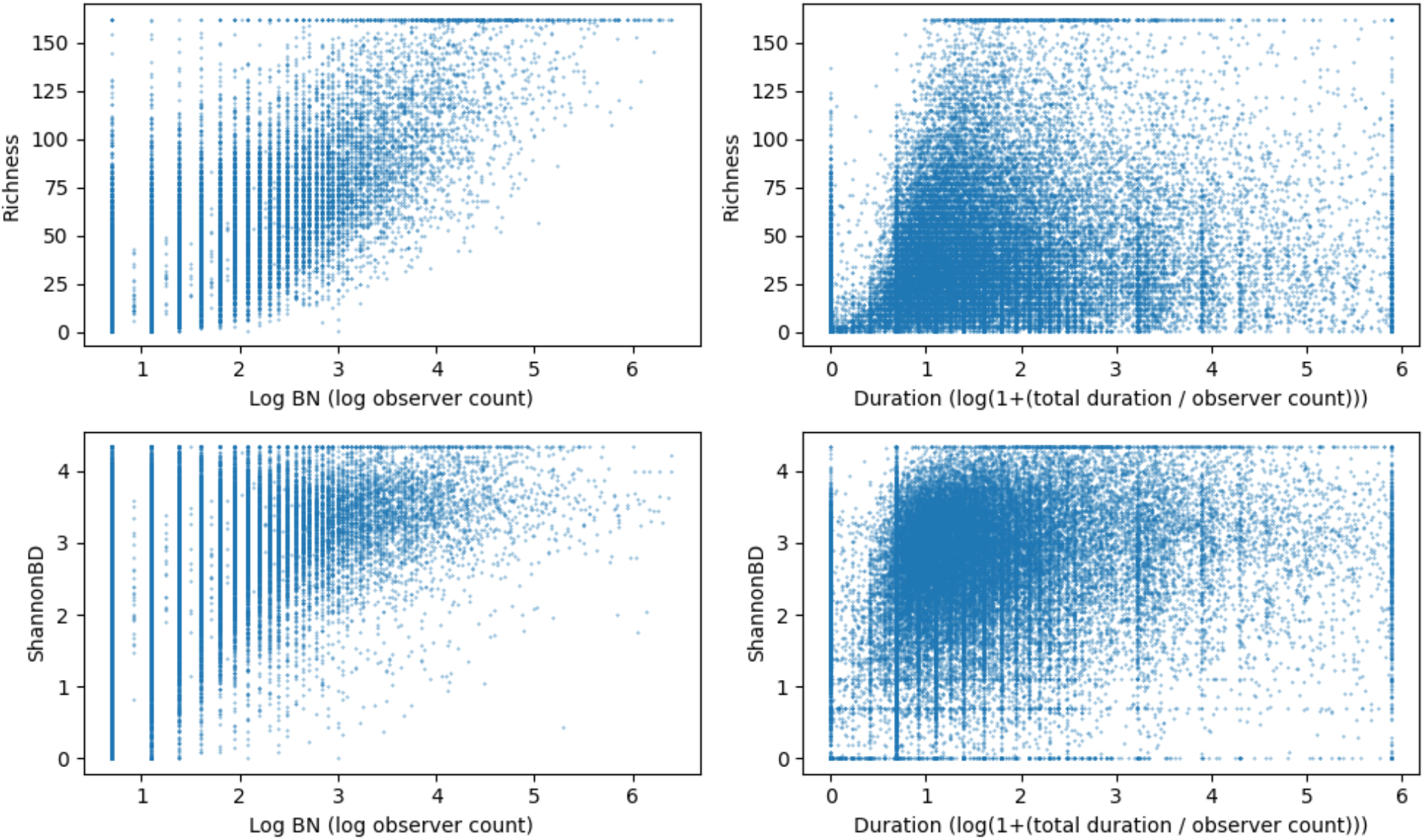
Relationships between sampling effort measures and observed bird diversity. Each point represents a county-month observation from 2014 to 2023. Richness (top row) and observed Shannon diversity (bottom row) show clearer positive relationships with log-transformed observer count (left) than with the study’s log-transformed duration-per-observer measure (right), which exhibits substantial dispersion. Observer count is transformed as log(1 + BN), and duration per observer as log[1 + (BT/BN)], where BN is observer count and BT is total recorded observation time. Horizontal concentrations at the upper bounds reflect winsorization of the diversity outcomes (as in the original study).

**Figure 3.**
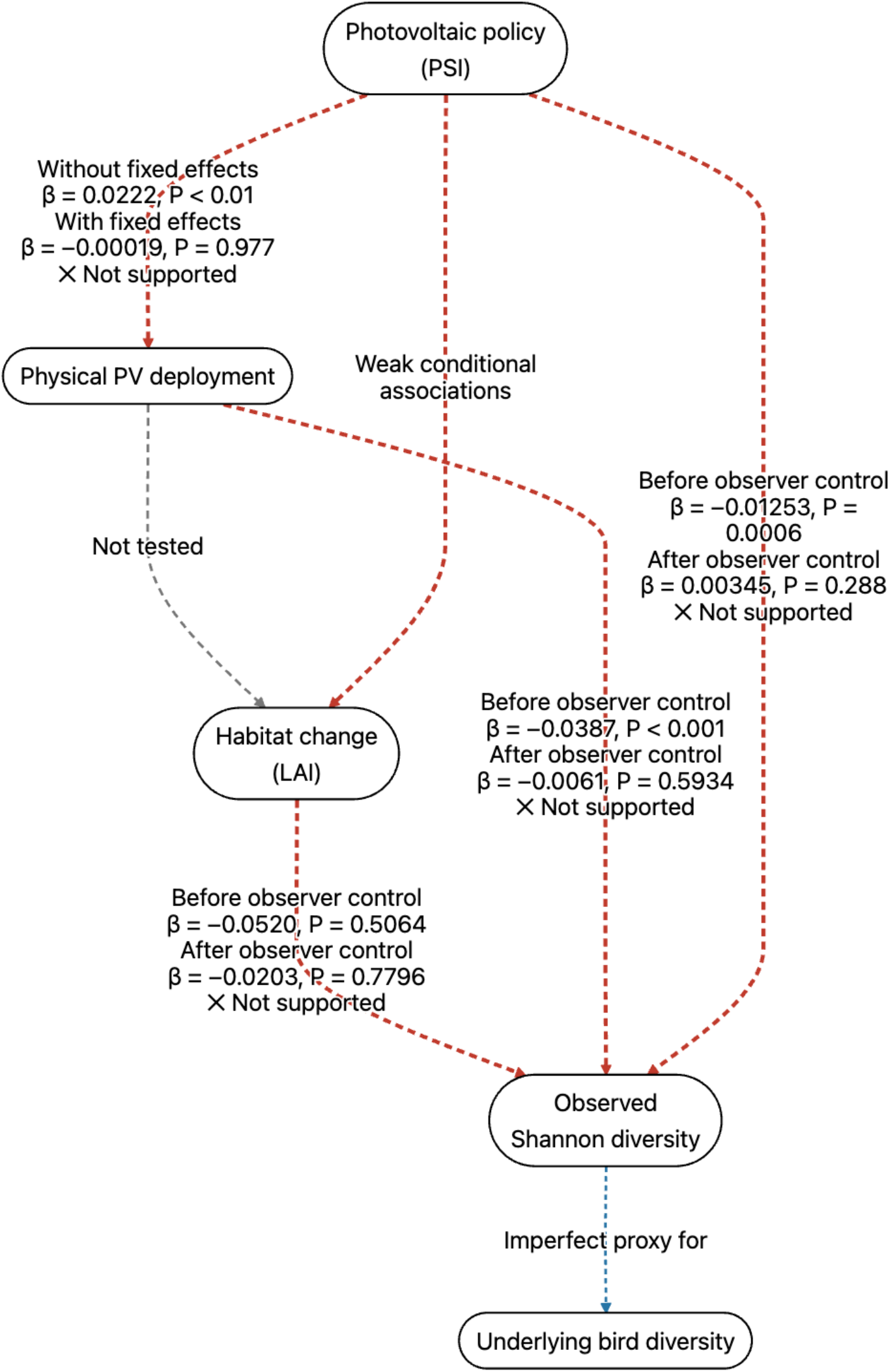
Evaluation of the proposed mechanistic pathway linking photovoltaic policy to bird diversity. The policy-to-deployment association disappeared after including county and year-month fixed effects. Associations of PSI, PV, and LAI with observed Shannon diversity became statistically indistinguishable from zero after controlling for observer count. The LAI– ShannonBD relationship was non-significant even before controlling for observer count. The direct PSI-LAI associations were weak. Red dashed paths indicate unsupported or weakly supported links, the gray path indicates an untested link, and the blue path emphasizes that observed Shannon diversity is an imperfect proxy for latent bird diversity.

**A further concern is the pronounced sampling domain shift over time** (*6, 7*). The number of recorded county-months also increased from 579 in 2014 to 13,719 in 2023, while the median observer count rose from one to three. Meanwhile, as reported earlier, the share of records with zero reported duration fell from 57.2% in 2014 and 68.1% in 2015 to 0.17% in 2023, and median total recorded time increased from zero to approximately seven to eight hours. These pronounced changes indicate a substantial temporal shift in participation and duration reporting rather than a stable sampling process. Other sampling factors, such as observers’ behavior and spatial sampling bias, may also change over time. The same sampling parameters may thus result in different observed biodiversity across years. Such sampling domain shift can have significant effect on trend estimate and needs to be controlled by statistical methods (*5, 7*).

Overall, the citizen science data indicate strong sampling bias that was unaccounted for in the main analysis. The lack of original data or the preprocessing methods further undermines the authors’ main result that PSI is negatively associated with bird diversity, a conclusion that has already been significantly weakened when incorporating just one source of sampling bias, the observer count.

### Problematic data and erroneous data preprocessing

The county-month aggregates in the dataset also contain implausible or erroneous effort and observation values (see the analysis notebook on GitHub for detailed information). For example, a large proportion of county-months had zero recorded sampling duration despite positive observer counts. This pattern was consistent with our inspection of the China Bird Report platform, where many checklists, particularly from the early years, displayed identical start and end timestamps (e.g., 2015-07-17 00:00 to 2015-07-17 00:00). The proportion declined from 68.1% in 2015 to 0.2% in 2023. These records may represent historical observations entered retrospectively to build life lists without duration information; alternatively, effort reporting may have been less standardized during the platform’s early years, when users were still becoming familiar with it. Because their underlying sampling effort cannot be reconstructed from the released data, these records cannot be reliably adjusted for effort or placed on a common sampling scale with other observations, making their use in effort-controlled analyses difficult to justify. **Moreover, reported birding time per observer exceeded 24 hours per month in 4**,**495 county-months (9.19%), exceeded the total duration of a 31-day month in 558 county-months (1.14%), and reached 78**,**473.3 hours in a county-month with only one observer** (四川省, 雅安市, 荥经县, 201405 [Yingjing County, Ya’an City, Sichuan Province, May 2014]), indicating serious problems with data filtering, aggregation, and preprocessing. Furthermore, in 10.4% of county-months, the Shannon and Simpson indices attain their perfect evenness values, a pattern consistent with identical count values being recorded for all species–a known issue in poorly standardized citizen science checklists. The data also likely contain severe spatiotemporal biases, as only 16.5% of the total county-month combinations were being sampled, likely due to data shortage. More seriously, the checklists at China Bird Report collected before 2025 are mixtures of complete and incomplete checklists, since the platform did not ask the users to submit such checklist completeness information (confirmed based on personal communications). Those data therefore mix presence-only observations with complete checklists, which are problematic to pool because the released data provide no way to distinguish between them or resolve the resulting ambiguity. These results suggest that the preprocessing, quality control, and even the choice of data were insufficiently rigorous for diversity assessment.

Method-wise, the study also failed to control for several important dimensions of sampling effort. Paradoxically, the duration-per-observer ratio used as an effort control discarded information about total sampling scale:

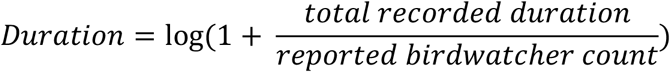

For example, one hour from one observer (e.g., data from 新疆维吾尔自治区 胡杨河市 202109 [Xinjiang province, Huyanghe county, Sep 2021]) and 49 hours from 49 observers (e.g., data from 安徽省 黄山市 歙县 202305 [Anhui province, Huangshan city, She county, May 2023]) both equal one, even though their sampling coverage can be very different. Both observer count and total observation time can affect opportunities to detect species and individuals. Therefore, when observations are pooled, the estimated richness and Shannon diversity can be much higher in 49-hour–49-observer cell than in 1-hour–1-observer cell. Controlling only for their ratio does not control for these dimensions of sampling effort. We are unaware of an established citizen science methodology that uses this ratio as the sole effort adjustment, and the paper provides neither a methodological reference nor a validation for doing so.

Ecological studies based on citizen science data also commonly retain checklist-level information so that duration, observer count, protocol, distance, completeness, and other sources of variation can be controlled for at the level at which observations were collected [e.g., ref. (*4, 8*)]. Pooling records into county-month averages makes this checklist-level variation untraceable and prevents direct standardization of heterogeneous sampling events. **Unfortunately, data from China Bird Report database are not publicly available for bulk download**. Due to this lack of original data and the unreported preprocessing of original data, the generating processes of those extreme and fractional values in the released data are unknown and unreproducible. We also cannot tell whether the pooled checklists were sampled once per group observation event, or were they being double counted. It is also impossible to determine whether the total number of observers consistently represents distinct observers, summed participant counts across checklists, or some mixture of these quantities.

### The reported R^2^ does not measure PSI’s explanatory power

The authors also misinterpreted their statistical results. The *R^2^* reported in Table 1 of the original paper is not PSI’s explanatory power; instead, the released code exports adjusted overall R^2^, including county and year-month effects. Our reanalysis reproduced *R^2^* = 0.2715 for the PSI– ShannonBD model [column 1, Table 1 in ref. (*1*)], but fixed effects alone contributed 0.2710.

After accounting for county and year-month fixed effects, PSI only explains 0.0733% of the variation remaining after those effects. In the fully controlled model [column 3, Table 1 in ref. (*1*)], its partial R^2^ is only 0.0478%, and its conditional increment is only 0.0941% of the model’s explained sum of squares. Similarly, the Table 2 in the original paper [ref. (*1*)] reported adjusted *R^2^* values near 0.992, which is also dominated by fixed effects (ranging from 0.991 to 0.992). After accounting for those effects, PSI’s partial *R^2^* for the regression against NDVI (column 1), nighttime (column 2), and leaf area (column 3) are 0.36%, 2.57% and 0.75%, respectively.

**Given these small explanatory powers, none of them is likely to be the strong mediating factors between PSI and diversity index**. The inferior greening mechanism therefore is unlikely to explain most of the changes in the observed bird diversity. It is worth noting we use partial *R^2^* only to correct the interpretation of the original model and the more consequential result is the sensitivity of the PSI coefficient to sampling effort controls. In ecology and conservation, statistical significance should not be interpreted as evidence of ecological importance, especially when sample size is large.

### No evidence for the proposed physical and mechanistic pathway

The policy–deployment physical pathway proposed by the authors is also unresolved. The authors argued that since PSI and PV area (the area of installed photovoltaic panels) are correlated, they provided a physical pathway of how PSI can influence bird diversity. Consistent with the paper, we reproduced the contemporaneous correlation between PSI and photovoltaic area (*r* = 0.0414), with the corresponding regression yielding *P* < 0.01. Although the authors emphasized its statistical significance, they did not discuss its magnitude: the correlation explains only approximately 0.17% of the variance and provides little empirical support for the proposed policy-to-deployment pathway. More importantly, although the main biodiversity models included county and year-month fixed effects, the authors did not apply the same specification when assessing this relationship. **When we included these fixed effects, PSI was no longer associated with photovoltaic area (*β*= −0.00019, *P* = 0.977)**. In the regression against ShannonBD, adding PV area leaves the PSI coefficient unchanged (*β*= -0.01254), meaning that PV and PSI share no upstream effect on ShannonBD in the two-way fixed effects setting. A potentially explanation for the lack of correlation is that the counties that have high PSI are also counties with low PV area and characterized by rolling hills, uneven terrains, and rainy summers. One possible explanation for the weak PSI–PV association is compensatory policy targeting. Counties with hilly terrain or rainier climates (e.g., Zhejiang, Hunan, and Guangxi provinces/autonomous region) may naturally attract less PV investment and therefore have less existing PV area [comparing Fig. 1A and 1B in ref. (*1*)]. Officials may respond with stronger policy support because these counties lag behind national targets, allowing high PSI to coexist with limited deployment; alternatively, the PV area data might be inaccurate. In conclusion, these results show no empirical connection between the policy index and contemporaneous installation area.

Additionally, the mechanistic pathways argued by the authors also do not hold. Results of fixed effect models showed that LAI is not a significant predictor of ShannonBD, either before controlling for observer count (*P* = 0.5064) or after controlling for observer count (*P* = 0.7796). Therefore, the mechanistic pathway from habitat changes to biological response is also not supported.

## Discussion

Our findings show that the negative association between photovoltaic policy and bird diversity in China reported by Zhang et al. (*1*) is highly sensitive to a basic sampling effort control. As a directly observable consequence, the county-level ShannonBD values plotted in Fig. 1C of Zhang et al. (*1*) show dramatic variations of diversity among spatially adjacent counties, which is implausible in ecological context. PSI predicts observer count, total observation time, and sample inclusion; once observer count is included, the main estimates and the instrumental variable estimate similarly become nonsignificant. The released data therefore cannot distinguish ecological change from changes in citizen science sampling effort. Yet observer count is only one of many sources of observation bias that the study failed to control. **This inability to separate ecological signals from the observation process is the central problem with the study, which invalidates their causal inference**.

The causal pathway proposed by Zhang et al. (*1*) depends on a chain from policy to physical deployment, from deployment to habitat change, and from habitat change to biological response. **None of these links is adequately supported by the released data, after controlling for observer count (Fig. 3)**. We are not arguing that photovoltaic development has no effect on birds. Rather, we are arguing that this study, as released, does not provide credible evidence for such an effect because the key explanatory variable is entangled with the process that generated the outcome.

Interestingly, one might argue that observer count could, in principle, act as a collider if both PSI and underlying bird diversity influenced observer participation and therefore should not be conditioned on. However, the county and year-month fixed effects identify the reported association primarily from changes in PSI and observed diversity within counties over time. For collider bias to explain the result, birdwatchers would need to detect and systematically adjust their participation in response to within-county diversity changes corresponding to only a 2.1% change in the Shannon index, which seems implausible. Meanwhile, the fixed effects do not account for temporal sampling domain shift within counties, and observer count directly affects the diversity recorded in citizen science data. These biases are impactful in temporal trend estimates and should not be left uncontrolled (*5, 7*).

**Understanding the ecological consequences of infrastructure development, including renewable-energy development, is essential for balancing decarbonization with biodiversity conservation** (*9*). Empirical studies have documented context-dependent effects of photovoltaic installations on birds. At a 180-ha photovoltaic facility in South Africa, bird richness and density were lower and community composition differed from adjacent untransformed habitat (*10*). Surveys of water-surface photovoltaic systems in China similarly found lower waterbird richness and density, generally lower Shannon diversity, and no observed nesting around photovoltaic installations (*11*). Conversely, some small photovoltaic farms established in intensively managed agricultural landscapes supported greater observed bird diversity than surrounding cropland (*12*). These studies suggest that outcomes of PV on bird diversity vary by environmental context, previous land use, facility design, and management.

Studies addressing this question can influence public understanding, conservation priorities, activities of nongovernmental organizations, and government policy. Therefore, the conclusions should be grounded in reliable data and rigorous statistical methods. Resolving the question of how solar expansion affects birds in China will require high-resolution data and proper study designs, either experimental or observational (*4, 13, 14*). Citizen science can substantially extend the geographic, temporal, and taxonomic coverage of biodiversity monitoring beyond what professional surveys alone can feasibly provide (*3*). Its value, however, depends on recognizing that the observation process is part of the data generating process. Where, when, and how people observe; how long and how far they travel; how many observers participate; their experience and reporting behavior; and whether they submit complete checklists all influence the species and individuals recorded (*4, 15, 16*). These factors can also change over time and differ among locations, producing sampling domain shifts that may resemble ecological trends (*5–7*). Analyses should therefore retain checklist-level information whenever possible, model effort variables separately, account for spatial and temporal sampling heterogeneity, and use suitable statistical methods and models for standardization of observation efforts [e.g. (*5, 17, 18*)]. Uncertainty arising from incomplete coverage, observer variability, and data processing should also be explicitly quantified and propagated into the final inference (*15*).

**These analyses also show that a claimed causal inference in a large dataset can erroneously create the appearance of ecological importance**. This understanding is especially important in large datasets, where small associations can be estimated precisely and therefore appear highly significant or have large effect sizes. However, without a deep understanding of the generation process and features of the ecological data, one can easily miss important variables or apply inappropriate analytical methods. Ecological interpretation should consequently consider robustness to alternative specifications, other potential confounders and mediators, and whether the proposed mechanism is supported by the data. **To properly understand the ecological data they are using, researchers from other fields (e.g**., **economics and geography) should collaborate closely with experienced ecologists to draw more reliable inferences and interpretations**. Statistical results alone, even when claimed to support causal conclusions, should be treated with particular caution when used to construct an elaborate ecological or policy narrative.

## Acknowledgments

We are grateful to researchers who independently reanalyzed the data and shared results that corroborated our findings. We thank the many ecologists, economists, birdwatchers, others who expressed interest in this work, shared insights and information, and offered encouragement and emotional support.

## Funding

No specific funding supported this work.

## Author contributions

Y.C. conceptualized the study, conducted the investigation and analyses, prepared the visualizations, and drafted the original manuscript. All authors contributed to reviewing and editing the manuscript.

## Competing interests

The authors declare no competing interests.

## Data and materials availability

The analyses use the data and code released by Zhang et al. at https://github.com/jianke22/China-s-solar-expansion-policy-reduces-bird-diversity. Reanalysis scripts and output tables are available at https://github.com/chenyangkang/reanalysis-Science-PSI-ShannonBD.

## AI-assisted drafting disclosure

OpenAI ChatGPT was used to assist with formatting of the code. The authors reviewed and verified the code, analyses and text and take full responsibility for the manuscript.

## Notes

https://github.com/chenyangkang/reanalysis-Science-PSI-ShannonBD

https://www.science.org/action/submitComment

### Competing Interest Statement

The authors have declared no competing interest.

